# Direct Supercritical Angle Localization Microscopy for Nanometer 3D Superresolution

**DOI:** 10.1101/2020.06.25.171058

**Authors:** Anindita Dasgupta, Joran Deschamps, Ulf Matti, Uwe Hübner, Jan Becker, Sebastian Strauss, Ralf Jungmann, Rainer Heintzmann, Jonas Ries

## Abstract

3D single molecule localization microscopy (SMLM) is an emerging superresolution method for structural cell biology, as it allows probing precise positions of proteins in cellular structures. Supercritical angle fluorescence strongly depends on the z-position of the fluorophore and can be used for z localization in a method called supercritical angle localization microscopy (SALM). Here, we realize the full potential of SALM by directly splitting supercritical and undercritical emission, using an ultra-high NA objective, and applying new fitting routines to extract precise intensities of single emitters, resulting in a four-fold improved z-resolution compared to the state of the art. We demonstrate nanometer isotropic localization precision on DNA origami structures, and on clathrin coated vesicles and microtubules in cells, illustrating the potential of SALM for cell biology.

---

For many biological questions the 3D organization of proteins is of high interest, therefore SMLM^1–3^ has been extended early on to go beyond 2D projections and to measure the 3D coordinates of single fluorescent emitters. This is most commonly achieved either by imaging fluorophores simultaneously in two or more focal planes^4,5^ or by introducing aberrations, such as astigmatism^6^, which allows extracting the z position from the shape of the point-spread function (PSF). Both approaches are robust and easy to implement, but their resolution is typically 3-fold worse in *z* and 1.5-fold worse in *x* and *y* compared to the lateral resolution of 2D SMLM^7,8^. Interferometric approaches such as iPALM^9^ or 4Pi-SMS^10,11^, can achieve a superb isotropic resolution, but are complicated to build and to use.

Another conceptually very different approach to determine z-coordinates of single fluorophores relies on detecting their supercritical angle fluorescence (SAF, Fig. 1a). SAF is a near-field effect that occurs if a fluorophore is in the vicinity of an interface with a higher refractive index (e.g. the microscope coverslip) and is near-exponentially decaying with the distance from the interface on the scale of 100 nm (Fig. 1b)^12,13^. Due to the blinking and the randomness of activation and de-activation of fluorophores in SMLM, their absolute intensities are not well defined. In order to reliably extract their z-positions, the under-critical angle fluorescence (UAF) has to be collected simultaneously and used for normalization. A major advantage of this approach is that the extracted z-coordinates report the absolute distance of the fluorophore from the coverslip.

**Figure 1:**
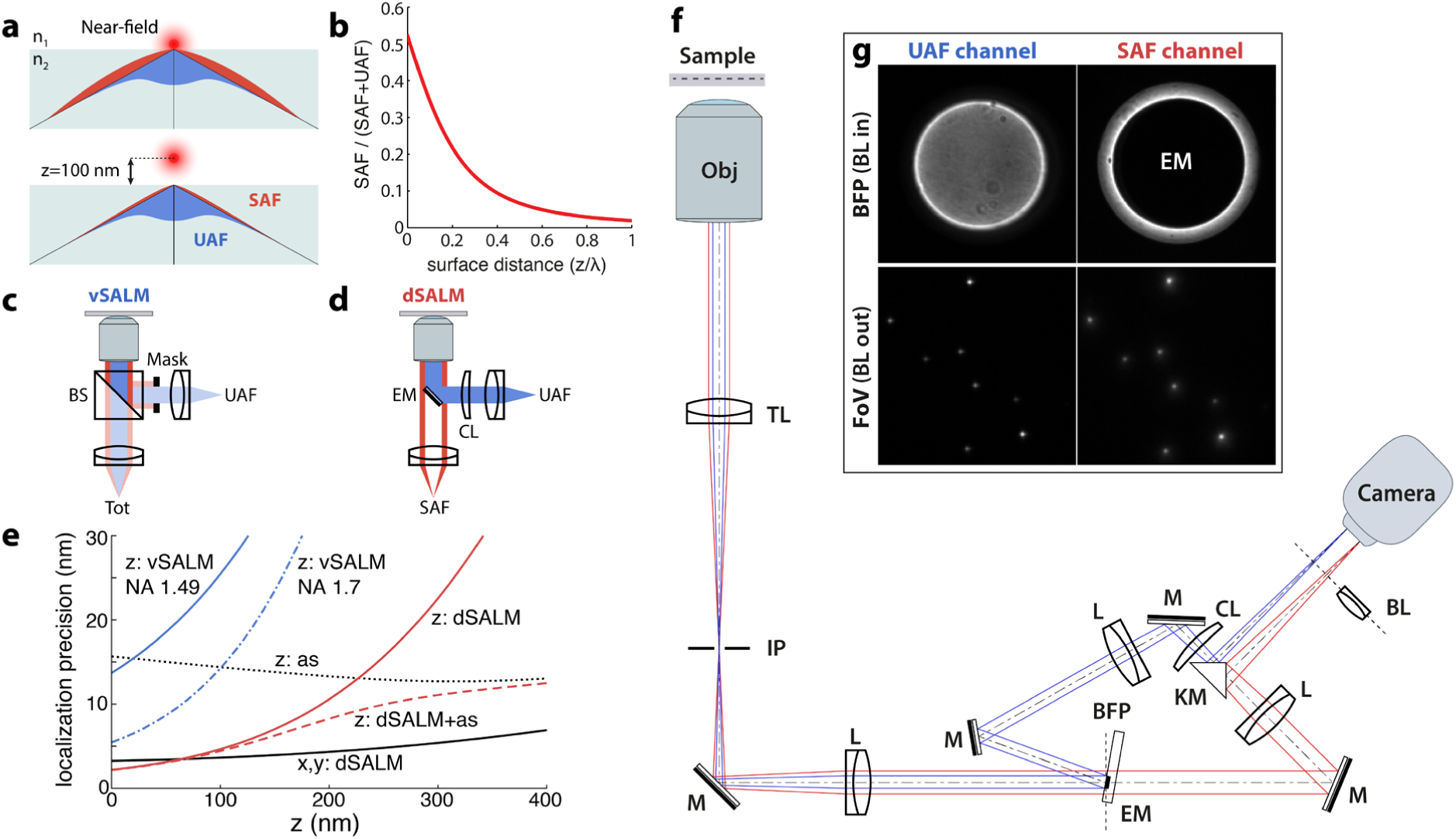
Concept of Super-critical Angle Localization Microscopy (SALM). **a**, Supercritical angle fluorescence (SAF) occurs when a fluorophore is close to a water-glass interface. **b**, SAF depends strongly on the fluorophore’s z position. **c**, virtual SALM (vSALM) splits the emission into two equal parts with a beam splitter (BS) and blocks out SAF with a mask in one channel. The z-position of the fluorophore is calculated from the ratio of total fluorescence to undercritical angle fluorescence (UAF). **d**, direct SALM (dSALM) splits SAF and UAF with an elliptical mirror (EM). An additional cylindrical lens (CL) in the UAF channel introduces weak astigmatism (as). **e**, theoretical localization precision in the axial direction calculated for an NA 1.49 objective in the vSALM configuration as used in ref ^14,15^ (blue line), and for an NA 1.70 objective for vSALM (dotted blue line) and dSALM (red line), astigmatism (dotted black line) and weighted average of dSALM and astigmatism (red dashed line). The solid black line indicates the lateral localization precision of dSALM. Calculations are based on experimentally derived PSFs and assume 5000 photons detected in the UAF channel, and a background of 50 photons per pixel in the UAF channel and 5 photons in the SAF channel. **f**, beam path for the dSALM implementation splitting SAF (red beam) and UAF (blue beam) before forming an image of the sample on the camera. BFP: back focal plane, BL: Bertrand lens, CL: cylindrical lens, EM: elliptical mirror, IP: image plane, KM: knife-edge prism mirror, L: lens, M: mirror, Obj: objective, TL: tube lens. **g**, UAF and SAF channels as seen on the camera with the Bertrand lens (BL) inserted in the beam path (upper panel) or without (lower panel).

This idea was implemented in proof-of-concept studies as *Supercritical-Angle Localization Microscopy* (SALM)^14^ or *Direct Optical Nanoscopy with Axially Localized Detection* (DONALD)^15^ in the so-called *virtual SAF* configuration^16^, i.e. by splitting the emission with a 50:50 beam splitter before detecting the total (UAF+SAF) emission in one emission channel and the UAF in a second channel after blocking the SAF with a mask (Fig. 1c). In the following we will refer to this implementation as ‘virtual SALM’ or vSALM. Although, in theory, vSALM has a higher resolution than astigmatism-based 3D SMLM in the close vicinity of the coverslip (Fig. 1e), experiments on biological samples failed to demonstrate this^14,15,17^. To extend the axial range of vSALM, it has been combined with astigmatism by introducing a cylindrical lens in the UAF channel^18^. Interestingly, SAF has a strong impact on the shape of a defocused PSF. Thus, it can be exploited for precise 3D localization in a simple single-channel setup using an ultra-high NA objective^19^. However, this approach requires precise knowledge of the z-dependent PSF, which is not easy to calibrate in presence of aberrations. Additionally, the required defocusing leads to a loss in lateral resolution. Related approaches used the enhanced near-field emission at metal^20,21^ or dielectric^22^ interfaces to precisely localize biological structures axially.

Direct splitting of SAF and UAF in SALM (direct SALM or dSALM) promises a several-fold improved z resolution and useable depth of field above the coverslip compared to vSALM (Fig. 1e, **Supplementary Fig. 1**). Indeed, SAF intensity can be quantified much more precisely if detected by itself than as part of the total intensity (SAF + UAF, as in vSALM), in particular when SAF is weak at larger z positions (see **Methods, theory**). The reason lies in the fact that the main source of noise is shot noise, therefore the relative error in determining photon numbers *N* scales with *δN*/*N* ∼ *N*^−1/2^. Additionally, and in contrast to vSALM (where splitting of the fluorescence halves the UAF signal), the entire UAF signal is used for normalization in dSALM, further improving precision. However, direct splitting of SAF and UAF had not been realized in SMLM because blocking the UAF in the SAF channel results in strong diffraction patterns dominating the SAF PSF^23^ (**Supplementary Fig. 2**), yielding an increased PSF size and preventing a reliable measurement of single molecule intensities.

Here, we overcome these challenges and realize the full potential of SALM by combining a) direct measurement of SAF with greatly increased signal to noise ratio by splitting it from UAF with a custom elliptical mirror, b) use of an ultra-high NA objective to increase the SAF signal and to decrease the effect of diffraction on the SAF PSF and c) new data analysis approaches that allow for precise determination of UAF and SAF intensities even in presence of a complex PSF. We demonstrate on 3D DNA origami structures that dSALM can fulfill its potential in terms of 3D resolution and show on biological samples that a combination with astigmatism leads to a remarkable z resolution over an extended axial range.

In the following, we give a short overview of the dSALM implementation, details can be found in the **Methods** section. To measure SAF and UAF of single fluorophores, we split the fluorescence in a plane conjugated to the objective back focal plane (BFP) with a custom-made elliptical mirror (**Supplementary Fig. 3**) and image both emission paths side-by-side on two halves of a camera (Fig. 1f). In the undercritical path, we insert a cylindrical lens to introduce weak astigmatism^18^. By fitting single molecules in the UAF channel with an experimentally derived PSF model^17^, we obtain their precise x and y coordinates and intensity, as well as less precise z coordinates (referred to as *z*_as_). Using these coordinates and a pre-determined transformation between the two channels, we apply a simple fit of an experimental SAF PSF model to the single molecule images in the SAF channel to extract their precise SAF intensities. From the ratio of SAF and UAF intensities we then calculate precise and absolute z positions, referred to as *z*_dSALM_, using the theoretical dependence of the SAF signal on z. For samples with an extended z-range, we calculate *z*_av_ as a weighted average of *z*_dSALM_ and *z*_as_^18^.

To validate the resolution of dSALM, we used DNA-PAINT^24^ to image DNA origami nanorulers that consisted of two rings of DNA binding sites separated by a distance of 30-nm (Fig. 2a-c). The two rings were easily resolved in side view reconstructions and a fit of z-profiles resulted in a standard deviation of z positions of 3.2 ± 1.4 nm and 5.8 ± 1.1 nm for the lower and upper ring, respectively (Fig. 2d). Due to possible deformations of the DNA structures and residual tilt, these are worst-case estimates for the experimental localization precision, the best-case estimate is the localization precision calculated from the Cramér-Rao Lower Bound (CRLB, **Methods**), which peaks at 2.7 nm (Fig. 2e).

**Figure 2:**
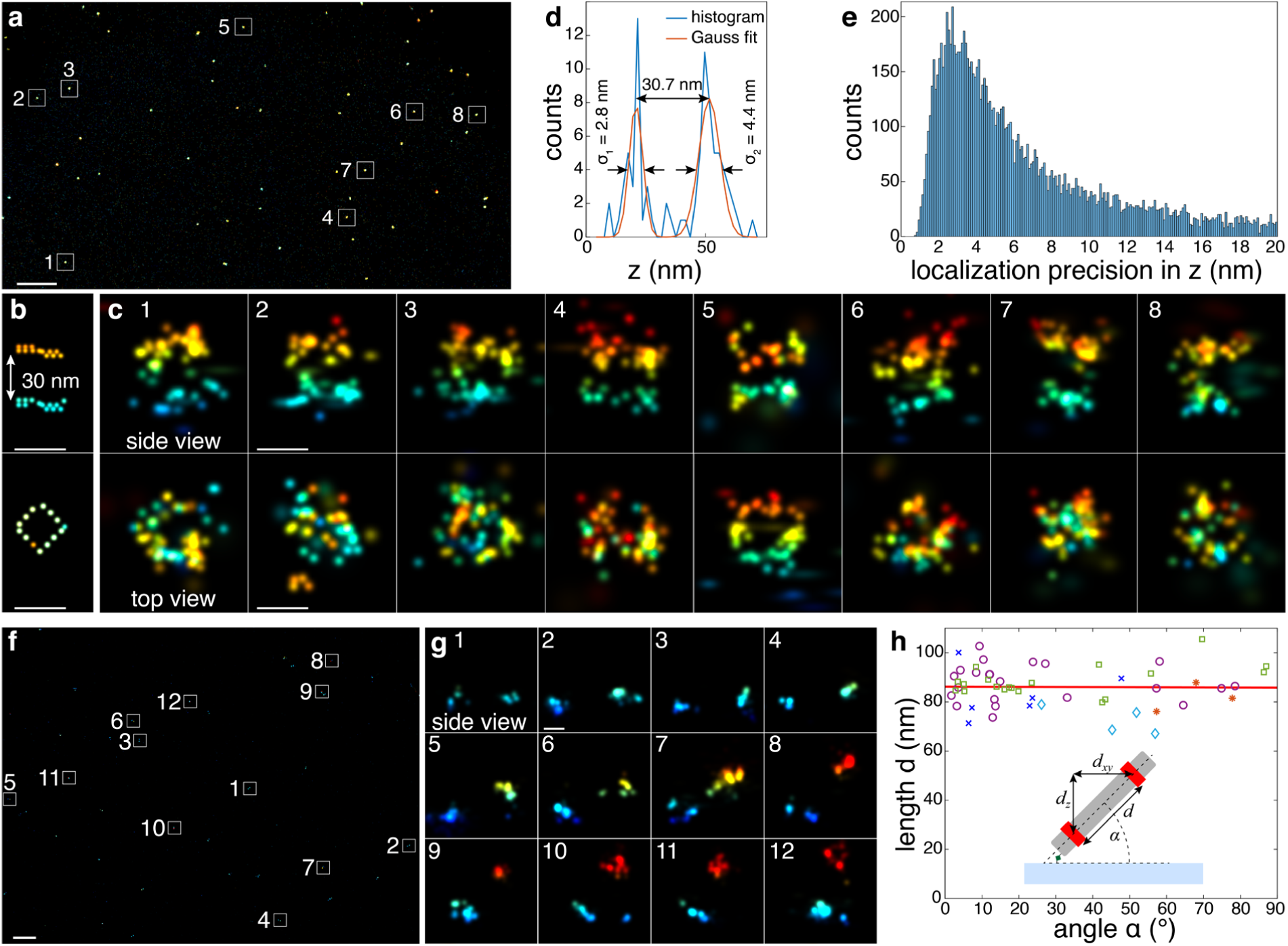
Validation. **a**, dSALM performed on 30-nm DNA origami rulers to test precision, overview image. **b**, schematic: each dot represents the position of a fluorophore binding site on the origami, as per design. **c**, side view and top view reconstructions of the rulers corresponding to the marked regions in a. Localizations are color-coded according to their z positions (*z*_SALM_). **d**, axial profile through the structure c7 with Gaussian fits resulting in a standard deviation of 2.8 nm for the lower and 4.4 nm of the upper ring. The average standard deviations of all structures c1-7 are 3.2 ± 1.4 nm and 5.8 ± 1.1 nm for the lower and upper ring, respectively, and their average distance is 27.6 ± 1.6 nm. **d**, the histogram of axial dSALM localization precisions (*δz*_dSALM_, **equation 7, Methods**) peaks at 2.7 nm. **f**, dSALM performed on 80-nm DNA origami rulers to test accuracy, overview image. **g**, side-view reconstructions corresponding to marked regions in f. **h**, measured length in dependence on the angle shows no depth-dependent deformations (Pearson correlation coefficient -0.015). Each symbol represents an independent experiment; red line is a linear fit to all data. Inset: sketch of the 80-nm DNA origami ruler. Scale bars: 1 µm (a, f), 30 nm (b, c, g).

In astigmatism-based SMLM, aberrations lead to a mismatch between model and data and consequently to systematic localization errors that depend on the distance of the fluorophore to the coverslip^25,26^. On the other hand, the intensity ratio between SAF and UAF is an absolute measure for the distance of the fluorophore from the coverslip and should be less sensitive to these aberrations. To test this, we imaged DNA origami rulers that consisted of two rings of DNA binding sites with a distance of 80-nm (Fig. 2f,g). Their immobilization on the coverslip resulted in a wide distribution of angles of the rulers with respect to the image plane. Any systematic localization errors would introduce a correlation between the angle with respect to the z-axis and the measured distance, something apparent in astigmatic 3D SMLM^25^. In our data (Fig. 2h), the measured distance did not depend on the angle, validating that dSALM allows for absolute z distance measurements with greatly reduced systematic errors.

To demonstrate the usefulness of our approach for biological research, we used speed-optimized DNA-PAINT^27^ to image two cellular structures, clathrin coated pits that form spherical assemblies with a diameter of around 150 nm (Fig. 3a,b, **Supplementary Fig. 4**), and microtubules, where the antibody labels form cylindrical arrangements with a diameter of approximately 50 nm around the filaments^28^ (Fig. 3c,d). These geometries are well visible in the side-view reconstructions (Fig. 3b,d). To extend the z range beyond the range accessible with dSALM, we based these reconstructions on *z*_av_. Compared to using *z*_dSALM_ alone, the z resolution away from the coverslip is improved (Fig. 3b1).

**Figure 3:**
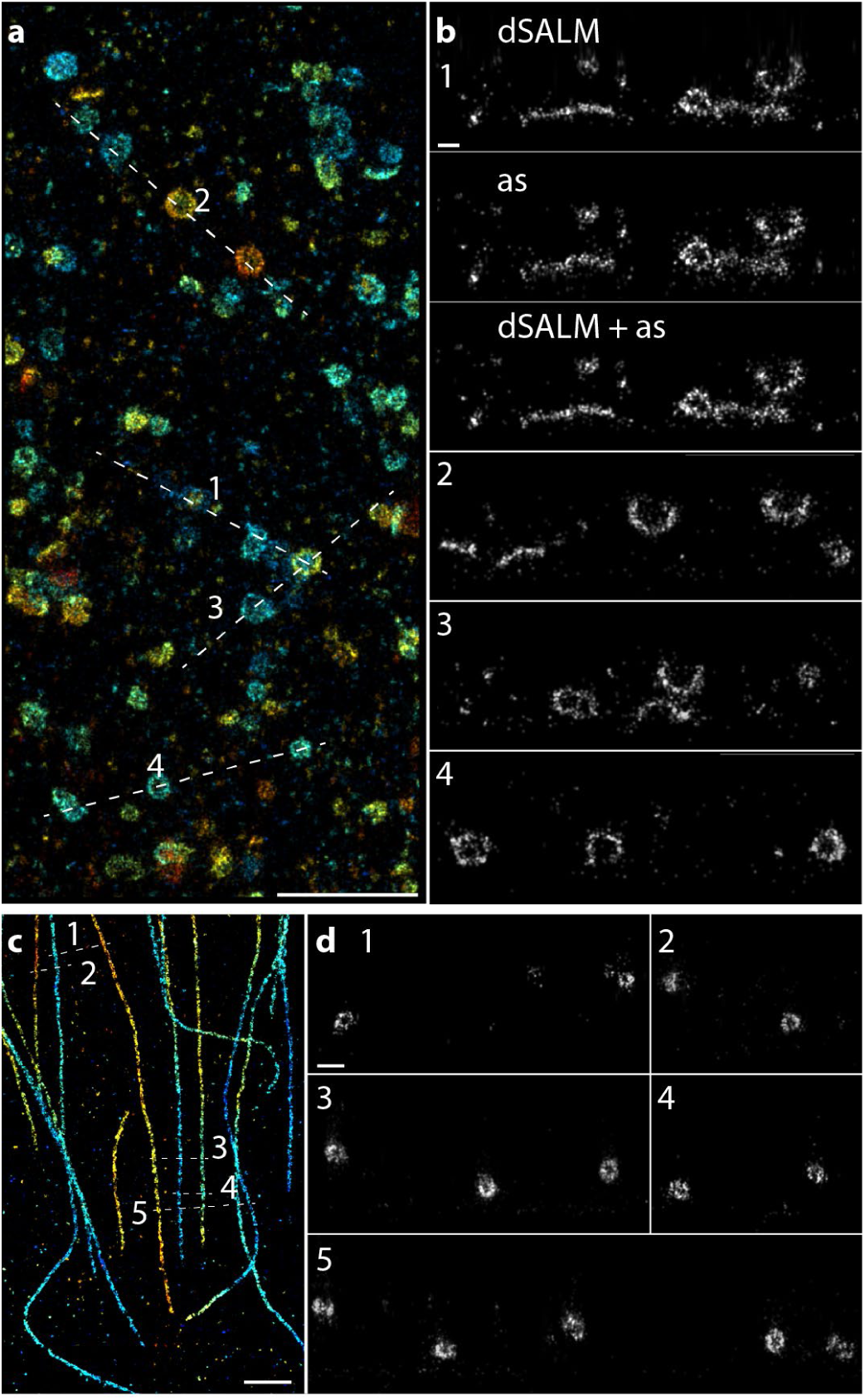
Application to biological structures. **a**, Clathrin coated pits, labeled with anti clathrin heavy and light chain primary and speed-optimized DNA-PAINT secondary antibodies. Localizations are color-coded according to their z positions. **b**, side-view reconstructions of regions as denoted in **a** (width of the line ROIs 50 nm). **b1** shows a comparison of a pure dSALM reconstruction (*z*_dSALM_), a reconstruction based on astigmatism (*z*_as_), and a combined dSALM + astigmatism reconstruction (*z*_av_), **b2-4** are based on *z*_av_. See **Supplementary Fig. 2** for additional reconstructions and localization precision histograms. **c**, Microtubules, labeled with anti alpha-tubulin primary and speed-optimized DNA-PAINT secondary antibodies. **d**, side-view reconstructions of regions as denoted in **c** (width of the line ROIs 200 nm). Scale bars: 1 µm (a,c), 100 nm (b,d).

To summarize, we developed dSALM, a 3D SMLM method that reaches a z resolution better than 10 nm near the coverslip, while retaining the high lateral resolution of 2D SMLM and providing absolute z distances from the coverslip. The resolution is comparable to that of interferometric SMLM, but with a much simpler and more robust setup. Compared to vSALM, a previous implementation of the method, the resolution and useable range above the coverslip are improved by more than 4-fold.

As SAF depends on the orientation of the transition dipole moment of the fluorophore, it should only be used with standard labeling approaches using fluorescent proteins, antibodies or self-labeling enzymes that allow for free rotation of the fluorophores. This is the case for DNA-PAINT, where the fluorophores are coupled to the imaging strand by a flexible linker and the docking strand contains unpaired nucleotides. The high refractive index immersion oil, required by the ultra-high NA objective, quickly degrades with blue or ultraviolet excitation, currently limiting the use of dSALM to SMLM with self-blinking dyes. Thus, the development of improved immersion oil is highly desirable to extend dSALM to photo-activatable fluorophores commonly used in SMLM. Finally, the circular field of view is limited to a diameter of 20 μm due to the strong field-dependent aberrations of the objective. A new generation of ultra-high NA objectives could increase the field of view and, thus, throughput. With these developments, dSALM has the potential to find widespread use in cell and structural biology and to enable new discoveries that are currently not accessible with standard SMLM.

## Acknowledgements

We thank Maurice Kahnwald and Aline Tschanz for their help with sample preparation. We would like to thank Alexander Jesacher for his help with the CRLB and PSF calculations and Mathieu Ribes and Alexander Jügler for help with production of the elliptical mirrors. This work was funded by the Deutsche Forschungs Gemeinschaft (DFG RI 2380/2), the European Research Council (CoG-724489 to J.R., StG-680241 to R.J.), the National Institutes of Health (U01 EB021223 to J.R.), the Human Frontier Science Program (RGY0065/2017 to J.R.), the Max Planck Society (R.J.) and the European Molecular Biology Laboratory (J.R., J.D., A.D., U.M.). S.S. acknowledges support by the QBM graduate school.

## Author contributions

J.R. and J.D. conceived the project. A.D., J.D. and J.R. built the microscope and performed and analyzed the measurements. U.M. prepared the samples. U.H., J.B. and R.H. produced the elliptical mirror. S.S. and R.J. provided the DNA-PAINT reagents. J.R., A.D., J.D. wrote the manuscript with input from all authors.

## METHODS

### Theory

#### Supercritical-angle fluorescence to extract z-positions of single fluorophores

A fluorophore close to a water-glass interface can couple its emission directly into the glass in an effect called surface-generated fluorescence or supercritical angle fluorescence^29,30^. This fluorescence strongly depends on the distance of the fluorophore from the interface. For freely rotating molecules, its intensity in the SAF channel *I*_*S*_(z) can be calculated numerically^29,31^:

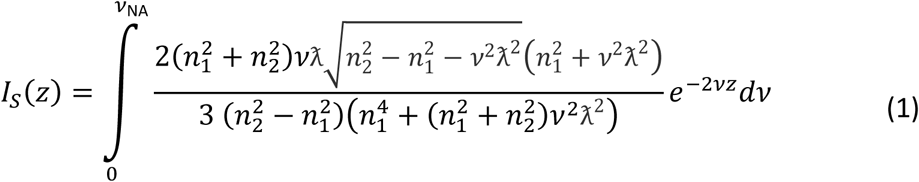

Here *λ* is the wavelength of the emitted light, 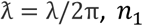 and *n*_2_ are the respective refractive indices of the buffer and the glass coverslip, 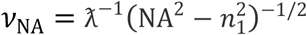, and the numerical aperture of the objective is defined as NA = *n*_2_ si*n* Θ_NA_.

The intensity in the UAF channel *I*_*U*_ is

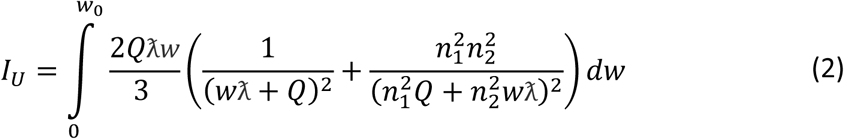

With 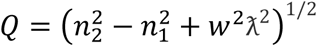 and 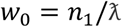.

In dSALM, we experimentally detect the number of photons emitted by a single fluorophore in the SAF channel as *N*_*S*_ and in the UAF channel as *N*_*U*_. To extract the z position of the fluorophore, we first calculate the theoretical intensity ratio from Eq (1) and (2):

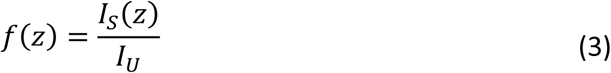

We then numerically invert this relationship and approximate *f*^−1^ = z(*f*) with a cubic spline. Then we can directly transform *N*_*S*_ and *N*_*U*_ to the fluorophore’s z position:

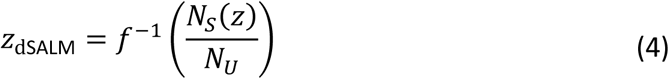

#### Calculating the Cramer-Rao Lower Bound (CRLB)

The CRLB is a best-case estimator of the precision of the fitting parameters. We use it to calculate the theoretical precision of dSALM and vSALM (Fig. 1e), and to assign experimental localization precisions to every single molecule. The CRLB can be calculated from the inverse of the Fisher Information matrix FI_*u,v*_ ^19,32^:

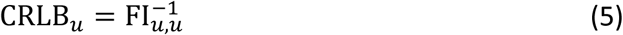

With

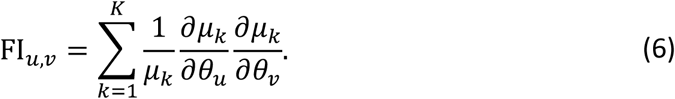

*μ*_*k*_(*θ*_*u*_) is the model describing the intensity in each pixel k. In our case, it is an experimentally derived spline-interpolated PSF model. *θ*_*u*_ = {*x, y, z*_*as*_, *N, b*} are the fitting parameters that include the position of the fluorophore *x, y, z*_as_, the number of photons *N* and the background per pixel *b*.

For **dSALM**, we estimate the lateral localization precisions 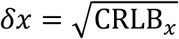 and 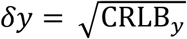 and the axial PSF-based localization precision 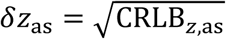 from the CRLB of the UAF channel only. The photometry-based axial localization precision in dSALM we calculate from the precision 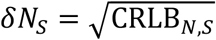 and 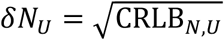 of the number of photons detected in each channel using eq. 5. To this end, we start with eq. 3, use the definition *f*(z) = *N*_*S*_/*N*_*U*_ and apply Gaussian error propagation^19^:

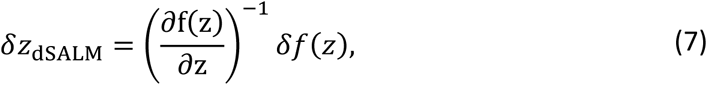

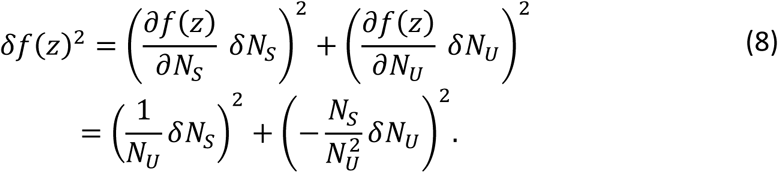

The combined axial localization precision from photometry *δZ*_Dsalm_ and PSF shape *δZ*_as_we calculate as the weighted average of each localization precision:

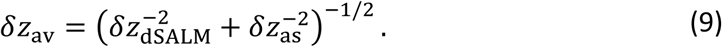

For **vSALM**, we estimated the lateral localization precision as the weighted average of the localization precisions of the UAF and total fluorescence channel, e.g.:

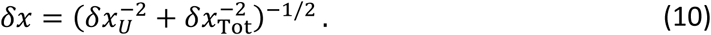

The axial localization precision based on photometry we calculate analogous to the dSALM example but with *N*_*S*_(*z*) replaced by *N*_Tot_(*z*) and *f*_*v*_(*z*) = *I*_Tot_(*z*)/*I*_*U*_ = *f*(*z*) + 1:

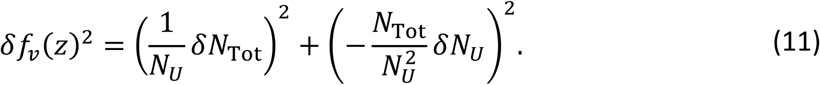

If we approximate *δN* with 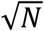, we can directly see where the improvement of dSALM vs vSALM comes from:

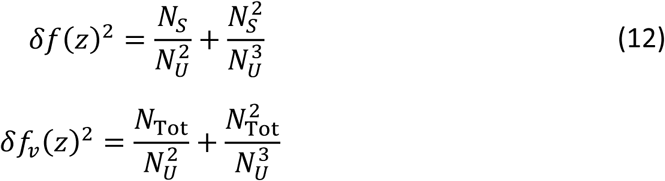

First, in vSALM the beam splitting leads to a decrease in *N*_*U*_ by a factor of 2 compared to dSALM. Second, especially for large z, *N*_Tot_≫ *N*_*S*_. Both increase *δf*_*v*_ compared to *δf*.

For calculating the CRLB in Fig. 1e, we assumed that we collect *N*_c_ = 5000 photons from a fluorophore far away from the coverslip. For fluorophores with a low quantum yield (as is the case for our experimental data with a quantum yield on the order of 30%) SAF competes not only with UAF, but mostly with the non-radiative decay. Thus, we made the approximation *N*_*U*_ = *N*_c_ and *N*_*S*_(*z*) = *f*(*z*)*N*_c_. For vSALM, after beam splitting *N*_*U*_ = *N*_c_ /2 and *N*_Tot_(z) = *N*_*U*_ + *N*_*S*_(z)/2. For fluorophores with high quantum yield, SAF competes with UAF and *N*_*U*_ + *N*_*S*_ = *N*_c_. Then *N*_*U*_ (*z*) = *N*_c_ /(1 + *f*(*z*)) and *N*_*S*_(*z*) = *N*_c_ *f*(*z*)/(1 + *f*(*z*)). The curves in Fig. 1e, although different in details, remain qualitatively the same.

### Microscope setup

#### Custom microscope

The detection beam path is shown in Fig. 1f. A laser combiner (iChromeMLE, Toptica) delivers the lasers (405 nm, 488 nm, 561 nm and 640 nm) to the microscope via a single-mode optical fiber. The laser beam is subsequently collimated by an achromatic lens (75 mm, Thorlabs) and circularly polarized by a quarter waveplate (Thorlabs). Another achromatic lens (dubbed illumination lens, 175 mm, Thorlabs), in 4f configuration with the objective, focuses the laser on the objective BFP. The fiber output is mounted on a linear stage (SmarAct) in order to switch the illumination mode between total internal reflection (TIR), highly inclined and laminated optical sheet (HiLo) or epi-illumination. Alternatively, a custom laser engine can be used to obtain homogeneous illumination, as described previously^33^. After the illumination lens, the lasers are reflected on a 4x dichroic mirror (F73-410, AHF) before the objective (NA 1.7 APON 100XHOTIRF, Olympus). The objective is housed in a z-stage (P-726, PI), while the sample position is controlled by a 2D stage (SmarAct). Fluorescence is collected by the objective and propagates through the 4x dichroic mirror. An intermediate image of the sample is formed by the tube lens (U-TLU, Olympus) outside of the microscope body. Another achromatic lens (200 mm, Thorlabs) in 4f configuration gives access to a plane conjugated with the objective BFP. In this plane, we inserted the elliptical mirror housed in a 2D translation mount (Thorlabs) at an angle of 10°, giving rise to the SAF and UAF channels. Finally, two achromatic lenses (250 mm, Thorlabs), in 4f configuration with the previous one, image the two channels side-by-side on two halves of an EMCCD camera (iXon 897, Andor). In the reflected channel (UAF), a 2 m cylindrical lens (SCX-50.8-1000.0-UV, CVI Laser Optics) is inserted to introduce weak astigmatism. Active axial drift correction is implemented by using a 785 nm laser (iBeamSmart, Toptica) coupled into the objective and reflected in TIR at the coverslip. The near infrared laser is then detected by a quadrant photo-diode (QP50-6 TO, First Sensor) and an analog signal from the sensor’s amplifier (LC-301DQD-PV, Laser Components) is directly fed to the objective z-stage controller. The microscope is entirely controlled by Micro-Manager^34^ using a custom plugin^35^.

#### Custom elliptical mirror

The mirror was produced by coating a layer of 65 nm protective Al_2_O_3_ onto 200 nm silver coating on a 5 mm thick AR-coated BK-7 glass (WG11050-A, Thorlabs) substrate with λ/10 nominal surface flatness. The layer structure was optimized aiming for a good compromise between predicted reflectivity (94.5% at 660 nm), material absorption and surface flatness. The optimization was performed with the help of a home-written computer program. The elliptical mirror used in this manuscript has a diameter of 5.50 mm × 5.58 mm, designed for a circular footprint when reflecting under an incident angle of 10°. The elliptical mirror was regularly cleaned under a microscope using an eyelash manipulator (commonly used for ultramicrotomy) to remove dust without damaging the surface.

### Sample preparation

#### Sample seeding

The ultra-high NA objective requires index-matching 20 mm round high refractive index coverslips (HIGHINDEX-CG, N4247800, Olympus). Since such coverslips are expensive, they were systematically cleaned and reused multiple times. To do so, coverslips were first cleaned gently with a tissue paper (Kimwipes, Kimtech) soaked in a 70 % EtOH solution. Next, they were kept overnight in 70% EtOH and finally air-dried for a few hours prior to the seeding of cells onto the coverslips. Then, cells (human bone osteosarcoma, U2OS) were seeded onto the coverslips 24 to 48 h before fixation in such a density that they reach a confluency of 50 to 70 % before further processing. Cells were grown in an incubation chamber providing 37 °C and 5% CO_2_ in growth medium (DMEM, catalog no. 11880-02, Gibco) containing 1× MEM NEAA (catalog no. 11140-035, Gibco), 1× GlutaMAX (catalog no. 35050-038, Gibco) and 10% (v/v) fetal bovine serum (catalog no. 10270-106, Gibco). Finally, shortly before fixation (see specific sample type description), coverslips were rinsed twice with warm PBS.

#### Imaging buffer for speed-optimized DNA-PAINT imaging

The following imaging buffer was used for imaging of microtubules and clathrin coated pits^27^. It comprised of 1x Trolox, 1x PCA, 1x PCD, 75 mM MgCl_2_, 5 mM Tris pH8, 1 mM EDTA, 0.05% Tween 20. Herein, 100x Trolox stock comprised of 100 mg Trolox in 430 µl methanol, 345 µl 1M NaOH, 3.2 ml H_2_O. 40x PCA stock comprised of 154 mg PCA in 10 ml H_2_O (adjusted to pH9.0 with 1M NaOH). 100x PCD stock comprised of 9.3 mg PCD in 13.3 ml (100 mM Tris pH8, 50 mM KCl, 1 mM EDTA, 50% Glycerol)

#### Beads on coverslip

Fluorescent beads were deposited on the coverslip by first preparing a solution with 360 µL water and 1 µl of fluorescent beads (0.1 µm TetraSpeck, T7279, Thermo Fisher Scientific). The solution was vortexed thoroughly. Meanwhile, 40 µl of 1M MgCl_2_ were pipetted onto the coverslip. Then, the beads solution was added to the coverslip and mixed. After 10 minutes the buffer was removed and 400 µl H_2_O was added to the coverslip.

#### 30-nm DNA origami preparation

In order to image the 30-nm 3D nanorulers (GATTA-PAINT 3D 30R ultimate line, GATTAquant), the high NA coverslips were washed three times with 500 μl of PBS. Then the coverslips were incubated with 200 μl of BSA-biotin solution (1 mg/ml in PBS) for 5 min. Upon removing the BSA--biotin solution, the coverslips were washed three times with 500 μl of PBS. Care was taken to ensure that the surface of the coverslips did not get scratched with the pipette tip. The coverslips were incubated with 200 μl of neutravidin solution (1 mg/ml in PBS) for 5 min. Upon removing the neutravidin solution the coverslips were washed three times with 500 μl of 1x PBS supplemented with immobilization buffer (1x IB: 10 mM MgCl_2_ + 500mM NaCl + 50mM Tris pH8). 7.5 μl of the 30 nm DNA origami solution was diluted with 200 μl 1x IB. The coverslips with the origami solution were incubated for 5 min. Eventually the coverslips were washed three times with 500 μl of 1x IB. The coverslips were then mounted into custom sample holders with 800 pM of imaging DNA strands coupled to Atto655 in imaging buffer comprised of 50 mM Tris pH 8, 10 mM MgCl_2_, 500 mM NaCl.

#### 80-nm DNA origami preparation

In the case of the 80-nm nanorulers (GATTA-PAINT 3D HiRes 80R, GATTAquant), the coverslips were treated following the same steps as described above for the 30-nm DNA Origami preparation. 5 μl of the 80 nm DNA origami solution was diluted with 100 μl 1x IB. The coverslips with the origami solution were incubated for 5 min. Eventually the coverslips were washed three times with 500 μl of 1x IB. The coverslips were then mounted into custom sample holders with 300 pM of imaging DNA strands coupled to Atto655 in imaging buffer comprised of 50 mM Tris pH 8, 10 mM MgCl_2_, 500 mM NaCl.

#### Indirect immunostaining of clathrin coated pit

Cells on coverslips were fixed at 37 °C with 3% formaldehyde (FA) in cytoskeleton buffer (10 mM MES, pH 6.2, 150 mM EGTA, 5 mM D-Glucose, 5 mM MgCl_2_) for 20 mins on a shaker. Then, the samples were rinsed once with 2 ml of freshly prepared 0.1% NaBH_4_ in PBS and then quenched in 0.1% NaBH_4_ in PBS for 7 min. After this, the samples were washed 3 times in PBS for 5 min each. In order to permeabilize the membranes, the samples were incubated in 100 µl of 0.01% Digitonin (Sigma D141) in PBS for 15 min. This was followed by 2 times washing in PBS for 5 min each. To block unspecific binding, the samples were placed in 100 µl ImageIT (I36933, Thermo Fisher Scientific) for 30 min and, following that, 30 min in 100 µl 2% BSA in PBS. For primary antibody labelling, 0.33 µl anti-clathrin heavy chain rabbit antibody (ab21679, Abcam) and 1 µl anti-clathrin light chain rabbit antibody (sc28276, Santa Cruz Biotechnology) were used in 100 µL 50% v/v ImageIT in PBS containing 2% (w/v) BSA overnight at 4 °C. Subsequently, binding of polyclonal donkey anti-rabbit secondary antibodies (catalog no. 711-005-152, Jackson ImmunoResearch) coupled to a DNA-PAINT docking site (TCCTCCTC) was achieved by placing the samples upside down onto a 1:100 dilution of the antibodies in PBS containing 2% (w/v) BSA overnight at 4 °C. The sample was imaged with in speed-optimized DNA-PAINT imaging buffer containing 17 pM of imager strands with the sequence AGGAGGA-ATTO643 (Eurofins Genomics).

#### Indirect immunostaining of microtubules

Coverslips containing U2OS were prefixed with 0.3% (w/v) glutaraldehyde (GA) in cytoskeleton buffer (CB: 10 mM MES pH6.1, 150 mM NaCl, 5 mM EGTA, 5 mM D-glucose, 5 mM MgCl_2_) containing 0.25% (v/v) Triton X-100 for 60 s before samples were incubated in 2% (w/v) GA in CB for 10 min. After briefly washing the samples with PBS, autofluorescence from GA fixation was quenched by incubation with freshly prepared PBS containing 0.1% (w/v) NaBH_4_ for 10 min. The sample was subsequently washed in PBS until no more bubbles were observed, usually 5x 5 min. Primary antibody staining was carried out by placing coverslips upside down onto a drop of primary antibody staining mix (mouse-anti-beta-tubulin, catalog no. T5293, Sigma-Aldrich, diluted 1:300 in PBS containing 2% (w/v) BSA) for 2-3h at RT. Weakly and unbound primary antibodies were washed off three times in PBS for 5 min each. Similarly, polyclonal donkey anti-mouse secondary antibodies (catalog no. 715-005-151, Jackson ImmunoResearch) coupled to a DNA-PAINT docking site (TCCTCCTC) was achieved by placing the samples upside down onto a 1:100 dilution of the antibodies in PBS containing 2% (w/v) BSA overnight at 4 °C. After washing thrice in PBS for 5 min each, post-fixation was carried out in 2.4% (v/v) formaldehyde in PBS for 30 min. Samples were quenched with 100 mM NH_4_Cl for 10 min, washed twice for 5 min in PBS and finally placed into a custom-made sample holder. The sample was imaged in speed-optimized DNA-PAINT imaging buffer containing 10 pM of imager with the sequence AGGAGGA-ATTO643^27^.

#### Antibody-DNA conjugation

Antibodies were conjugated to DNA-PAINT docking sites via DBCO-sulfo-NHS ester chemistry as previously reported^36^. In brief, antibodies were reacted with 20-fold excess of a bifunctional DBCO-sulfo-NHS ester (Jena Biosciences, cat: CLK-A124-10). Unreacted linker was removed using Zeba Spin Desalting columns (0.5 ml, 40k MWCO, Thermo Fisher Scientific, cat: 89882). Azide-DNA (C3-azide-TCCTCCTC, Metabion) was added to the DBCO-antibodies with a 10-fold molar excess and reacted overnight at 4°C. Afterwards, buffer was exchanged to PBS using Amicon centrifugal filters (100k MWCO).

### Data acquisition

#### Determining the region of low optical aberrations

The ultra-high NA objective from Olympus displays strong field-dependent aberrations. Only a small part at the center of the objective allowed for efficient PSF averaging and precise estimation of intensities. We estimated the position of this region by taking z-stacks on 10 different fields of view without EM gain, fromb-1.5 µm to +1.5 µm with 50 nm step size. We then fitted the z-stacks using an experimentally derived PSF model and calculated the displacement standard deviations in x and y for each bead. The analysis showed a strong dependency of the displacement standard deviation on the beads distance to the center of the objective. Based on these results, we limited all our experiments to a circle of radius 10 µm around the objective center.

#### Oil preparation

Before each series of experiments, 30 µl of high refractive index oil (Series M 1.780, Cargille) was centrifuged at 10000 rpm for 2 min, causing oil crystals to sediment. Only 10 µl of the supernatant is used at a time on the objective. When imaging nanorulers and biological samples, the oil was exchanged for every new region of interest.

#### Bead stacks

About 15 fields of view were manually chosen, concentrating on regions with few non-overlapping beads. The stacks were recorded at 30 ms exposure time, without EM gain from -1.2 µm to +1.2 µm around the focus in steps of 30-nm. A mean excitation intensity of 0.9 kW/cm^2^ was used.

#### DNA-PAINT nanorulers

The 80-nm nanorulers were imaged with an EM gain of 100 and an exposure time of 300 ms. The excitation was set so that we obtained a power density of about 9 kW/cm^2^ using two 640 nm laser diodes from the laser engine (homogeneous illumination). Imaging was stopped after 80,000 frames. The 30-nm nanorulers were recorded with an EM gain of 100 and 500 ms exposure time. Imaging was performed under total internal reflection illumination for 12,000 frames using the commercial laser combiner (640 nm laser) at a power density of 10 kW/cm^2^.

#### Biological samples

Images of Clathrin-mediated endocytosis were acquired with 100 ms exposure time and an EM gain of 100. We typically imaged the samples for 100,000 frames in HiLo (640 nm laser form the commercial combiner). The illumination led to an estimated power density of 6.5 kW/cm^2^. Microtubules images were acquired using the same settings, albeit with a slightly lower power density (5.6 kW/cm^2^).

### Data analysis

All data analysis was performed with a custom MATLAB software called Superresolution Microscopy Analysis Platform (SMAP, https://github.com/jries/SMAP).

#### Calibration of the experimental PSF model

In dSALM, the z position of a single fluorophore is extracted from its relative intensities in the supercritical and undercritical channel. This requires the precise measurement of these intensities in the camera images, which is challenging because diffraction causes the PSF to be very different from one channel to the other, and depends on z. Here, we overcome this challenge by fitting the fluorophore images with experimentally derived PSF models. To this end we use stacks of beads immobilized on a coverslip. We generated a spline-interpolated PSF model for each channel after registering and averaging about 40 beads following previous work^17^. In particular, we only considered beads within a circle of 10 µm radius around the center of the objective field of view, a region where the objective PSF does not display strong field-dependent aberrations. The resulting PSF calibration is used for analysis of all data.

#### Quantification of SAF and UAF intensities

For the nanorulers and the biological samples, the localization microscopy raw data was fitted with the experimental PSF models for the UAF and SAF channel, using maximum likelihood estimation^17^. From the fitted positions of fluorophores detected in both channels, we generated a projective transformation from the SAF to the UAF channel. Based on the 3D coordinates of every fluorophore in the UAF channel, the transformation and the PSF model for the supercritical channel, we then applied a second fitting step, in which we calculated the precise shape of the SAF PSF at the position of the fluorophore in the SAF channel from the SAF PSF model and use it to fit intensity and background. This leads to a much more precise intensity estimation than during the first fitting step and allows for precise intensity estimation for fluorophores that due to their z position were too weak to be detected in the SAF channel in the first fitting step.

#### Calculation of z positions

From the relative intensities of the single fluorophores in the SAF and UAF channel we calculated *z*_dSALM_ for every single-molecule blinking event according to eq. (4). The astigmatism induced in the UAF channel helps in the initial fitting step by eliminating the degeneracy of the unmodified 2D PSF^17^ and it extends the 3D resolution of dSALM beyond the range of SAF detection^18^. To combine *z*_dSALM_ with the z-position from the initial fitting of the experimental astigmatic PSF *z*_as_, precise knowledge of the coverslip position is required. As this position changes due to drift, we first perform a 3D drift correction based on redundant cross-correlation^17^. Then, we plot for all localizations their intensity ratio *f*^*ex*^ = *N*_*S*_/*N*_*U*_ vs the drift-corrected 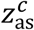. Finally, we determine the glass position PSF *Z*_as,0_ from a fit of *f*(*z* − *z*_as,0_) (eq 3) to these data points, treating *z*_as,0_ as the only free fitting parameter. Using the respective CRLB as weights, we then combine *z*_dSALM_ and 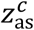 as a weighted average to *z*_av_:

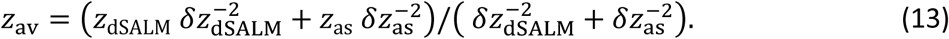

#### Post-processing and rendering

After fitting, localizations stemming from the same fluorophore and persisting over several frames were merged into a single localization. Localizations with a poor fit, resulting probably from overlapping localizations, were identified by their normalized log-likelihood value and filtered out (values lower than -2). In addition, dim localizations leading to a lateral localization precision of less than 10 nm were filtered out. Superresolution images were reconstructed by rendering each localization as a Gaussian with a size proportional to the localization precision.

#### Analysis of 30-nm DNA origami

Structures were manually segmented. As the DNA origami structures can be tilted with respect to the coverslip, they were manually rotated to obtain side views using the 3D viewer in SMAP. Histograms of the rotated z-position were fitted with a double Gaussian to extract the distance and the standard deviation of each ring.

#### Analysis of 80-nm DNA origami structures

We manually segmented 53 DNA origami nanopillars. We then used a DBSCAN algorithm^37^ to assign the localizations to clusters and adjusted the two parameters (size scale and minimum number of localizations in the neighborhood), resulting in two main clusters per structure. We calculated the weighted average of the x, y and z coordinates for each cluster. From the average positions, we calculated their distance in 3D and their projected distance in the x, y plane, and from these values the angle of the pillar with respect to the image plane.

---

**Supplementary Figure 1:**
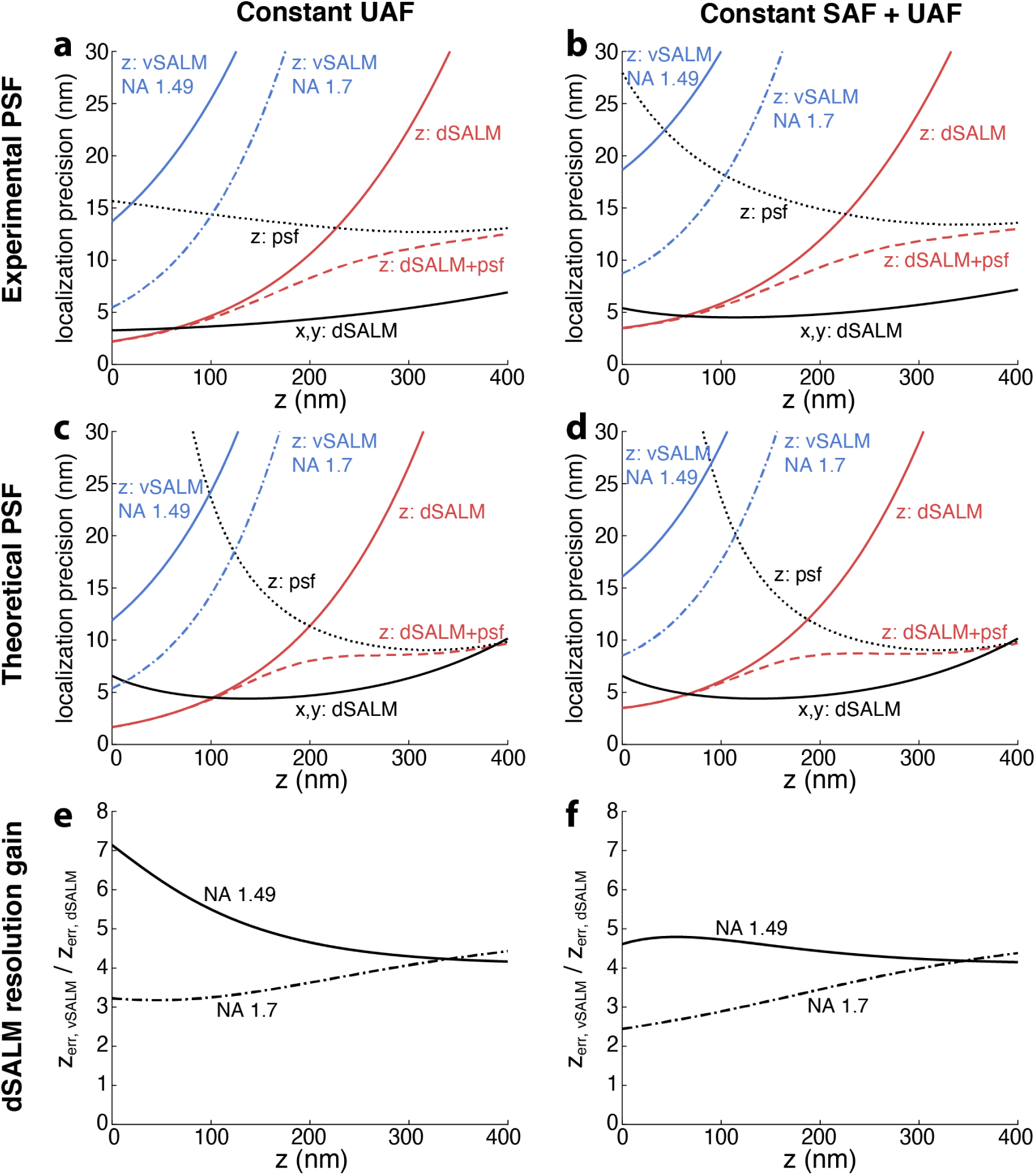
Theoretical resolution of supercritical-angle localization microscopy for vSALM with an NA 1.49 and NA 1.7 objective and dSALM with an NA 1.7 objective. psf denotes the z localization precision obtained from fitting the PSF shape as common for astigmatic SMLM. Comparison of experimentally derived PSF (a,b) and PSF calculated according to *Zelger et al*., *Biomed. Opt. Express 11 (2020)* (c,d). (e,f) display the ratio of z localization precisions between vSALM and dSALM, which corresponds to the relative gain in resolution of dSALM vs. vSALM. (a,c,e) assume constant UAF, as an approximation for fluorophores with a low quantum yield where SAF competes mostly with non-radiative decay. (b,d,f) assume constant total fluorescence, as an approximation for fluorophores with a high quantum yield.

**Supplementary Figure 2:**
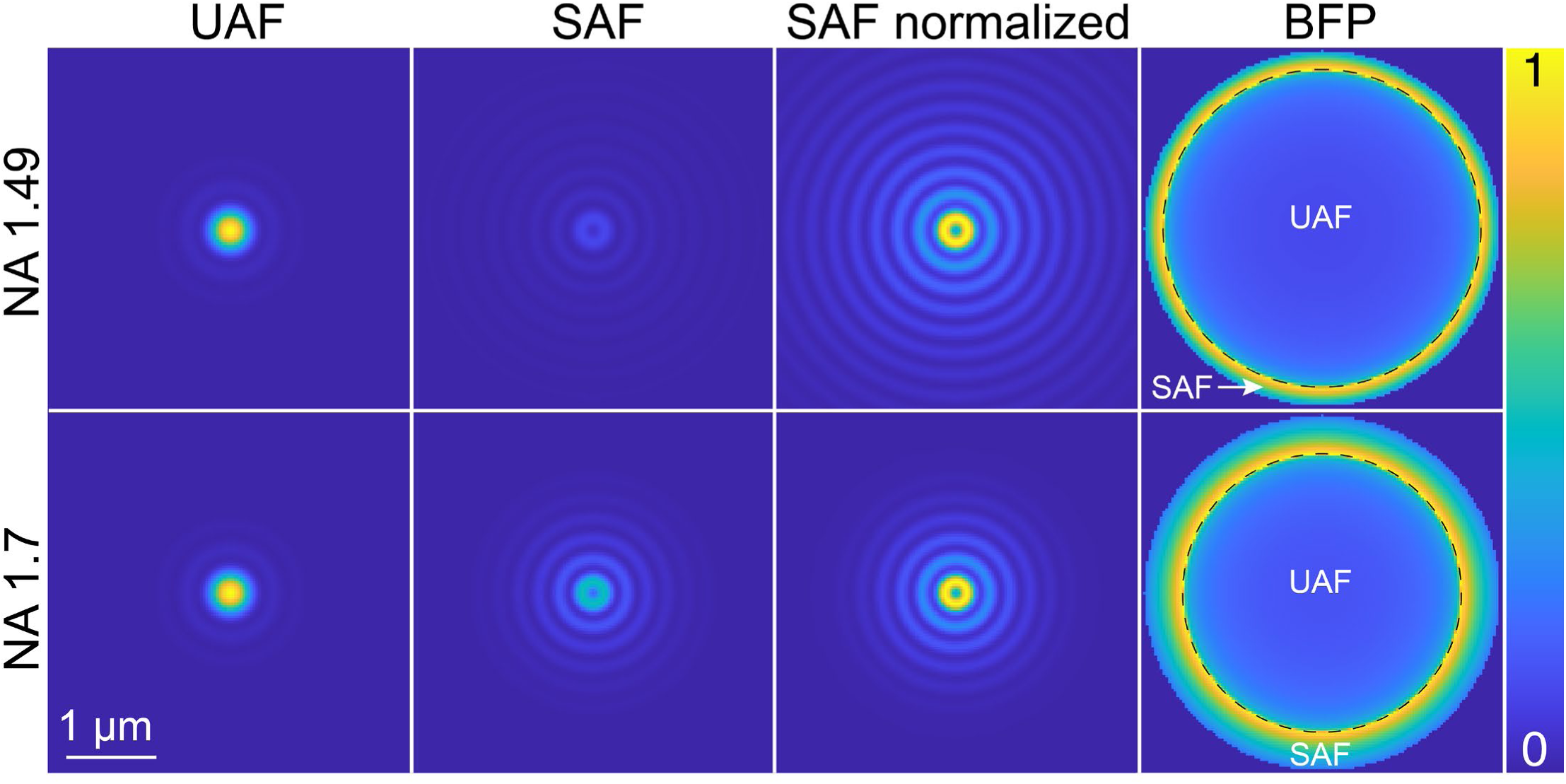
Calculated PSFs for the NA 1.49 and the NA 1.7 objective in the dSALM configuration for the UAF and SAF channel, and the corresponding back focal plane intensity distribution. The emitter is at the coverslip (z=0). Calculations based on code from *Zelger et al*., *Biomed. Opt. Express 11 (2020)*. The SAF PSF displays strong diffraction rings leading to a large PSF size, especially visible in the ‘SAF normalized’, where the intensity is normalized to its maximum. In the calculated BFP images, the mirror separating UAF and SAF is denoted with a dashed line.

**Supplementary Figure 3:**
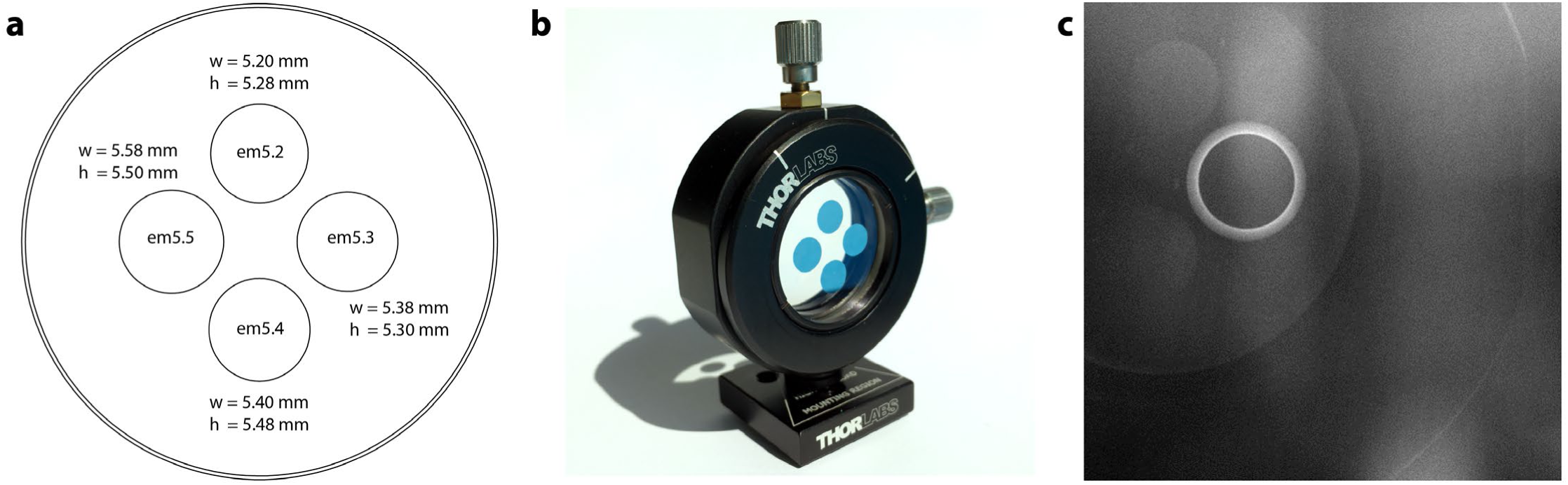
Elliptical masks. **a**, Specification of the masks dimension, designed to be placed at an angle of 10° in the beam path. **b**, Picture of the four elliptical masks deposited on the glass window and housed in a 2D translation mount. **c**, Camera image of the BFP with stray light highlighting the masks and the holder.

**Supplementary Figure 4:**
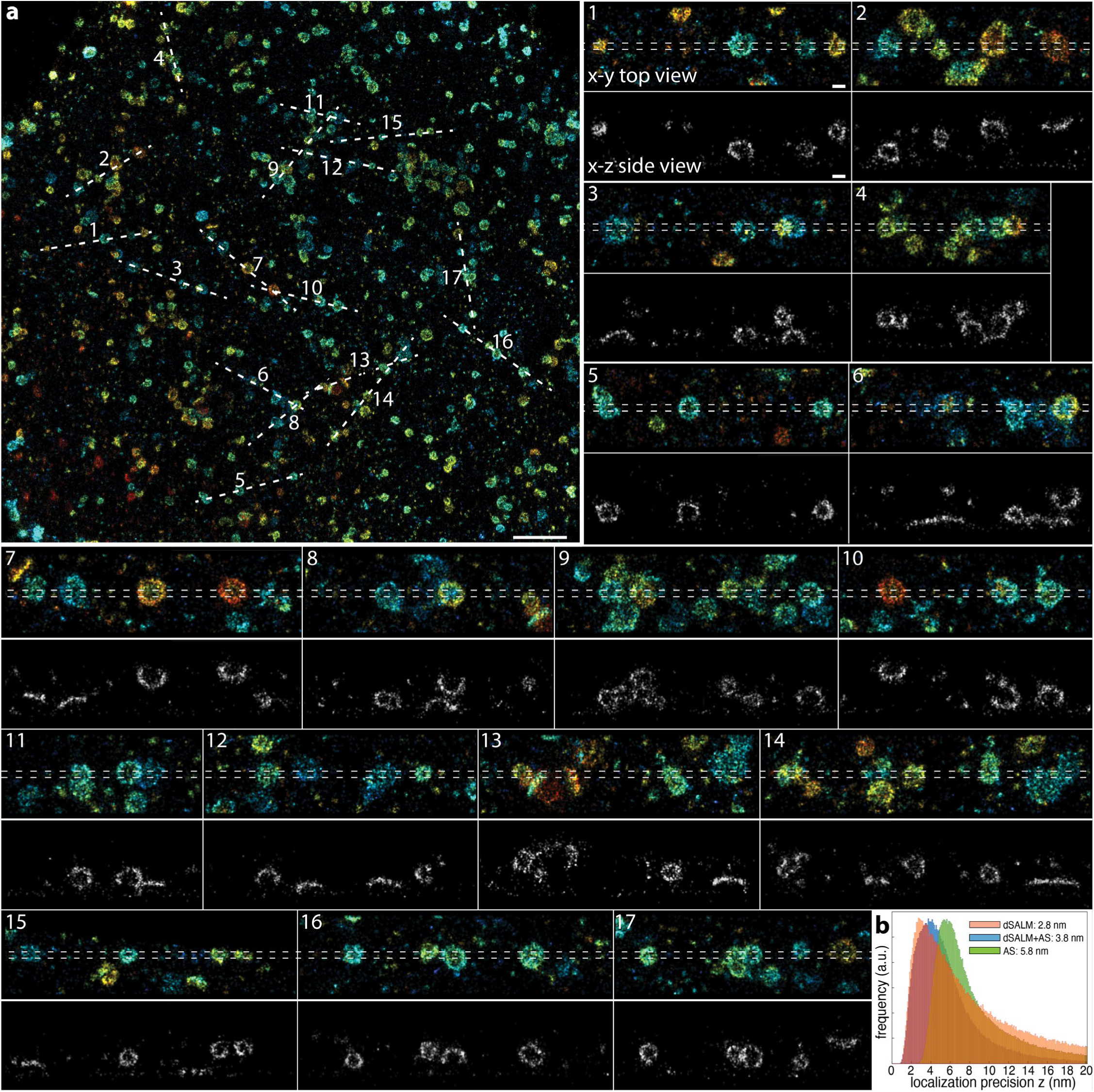
Clathrin coated pits. **a**, Clathrin coated pits, labeled with anti clathrin heavy and light chain primary and fast DNA-PAINT secondary antibodies. Localizations are color-coded according to their z positions. Insets represent the various sites of the overview image in x-y top view (top) and x-z side view (bottom). **b**, Histograms of the localization precision (CRLB) of the z-coordinate *z*_dSALM_ for dSALM (orange), *z*_av_ for combined dSALM and astigmatism (blue) and *z*_as_ for astigmatism alone (green). Scale bars 1 µm (a), 100 nm (1).

